# Optimal density of bacterial cells

**DOI:** 10.1101/2020.11.18.388744

**Authors:** Tin Yau Pang, Martin J. Lercher

## Abstract

A substantial fraction of the bacterial cytosol is occupied by catalysts and their substrates. While a higher volume density of catalysts and substrates might boost biochemical fluxes, the resulting molecular crowding can slow down diffusion, perturb the reactions’ Gibbs free energies, and reduce the catalytic efficiency of proteins. Due to these tradeoffs, dry mass density likely possesses an optimum that facilitates maximal cellular growth and that is interdependent on the cytosolic molecule size distribution. Here, we analyze the balanced growth of a model cell, accounting systematically for crowding effects on reaction kinetics. Its optimal cytosolic volume occupancy depends on the nutrient-dependent resource allocation into large ribosomal vs. small metabolic macromolecules, reflecting a tradeoff between the saturation of metabolic enzymes, favoring larger occupancies with higher encounter rates, and the inhibition of the ribosomes, favoring lower occupancies with unhindered diffusion of tRNAs. Our predictions across growth rates are quantitatively consistent with the experimentally observed reduction in volume occupancy on rich media compared to minimal media in *E. coli.* Strong deviations from optimal cytosolic occupancy only lead to minute reductions in growth rate, which are nevertheless evolutionarily relevant due to large bacterial population sizes. In sum, cytosolic density variation in bacterial cells appears to be consistent with an optimality principle of cellular efficiency.

## Introduction

The dry mass dissolved in the major compartment of bacterial cells, the cytosol, comprises hundreds of molecular species, including proteins, metabolites, polysaccharides, and nucleic acids. These molecules can be roughly classified into two sectors: the ribosomal sector, dominated by ribosomes and tRNA; and the non-ribosomal sector, comprising mostly metabolites, enzymes, and other proteins [1]. The molecules in these two sectors have very different size distributions: the ribosome is 65 times larger than the median enzyme size (2,600kDa [2] vs. 40kDa [3]), and tRNAs are about 300 times larger than typical metabolites (26kDa [4] vs. 89Da, the mass of alanine). The allocation of dry mass between the two sectors of the cellular economy can be summarized by a single parameter, the growth rate *μ*, with the dry mass fraction of the ribosomal sector increasing almost linearly with *μ* [1,5]. Accordingly, the ribosome-rich cytosol at fast growth in nutrient-rich environments and the ribosome-meager cytosol at slow growth in nutritionally poor environments exhibit very different distributions of molecule sizes.

Experiments have found an approximately constant dry mass density of the cytosol at 300g/L across minimal nutrient conditions with low growth rates, but a roughly 10% less dense cytosol in a rich medium supporting high growth rates [6]. We hypothesized that the observed difference in cytosolic density between slow and fast growth may be an evolutionary consequence of the differences in molecular composition. Cellular physiology has evolved under natural selection; thus, if the level of molecular crowding—the dry mass density of the cytosol—affects cellular efficiency and hence fitness, we expect density regulation to have evolved to a near-optimal, possibly condition-dependent state. While previous work has explored the influence of (macro-)molecular crowding on bacterial physiology and growth rates, these analyses assumed a constant, hard limit on the total cytosolic protein [7,8] or dry mass [9,10] concentration. These works do not justify the existence and magnitude of the density limit based on physicochemistry, and they cannot explain differences in the level of crowding (dry mass density) across conditions.

Biochemical reaction fluxes typically increase with increasing encounter rates of the molecule species involved; ignoring crowding effects on diffusion, encounter rates increase with increasing density of the respective molecules. At the same time, the molecular crowding caused by other molecule species in the background (volume-excluding co-solutes) affects fluxes in at least three distinct ways. (i) Crowding slows down the diffusion of a catalyst and its substrates, thereby reducing their encounter rates [11,12]. (ii) Crowding limits the available volume and thus reduces a solute’s entropy (the “excluded volume phenomenon”), thereby changing the free energies of the molecules involved in the reaction and consequently shifting the equilibrium concentrations of substrates and products [13]. (iii) Crowding can affect the structure of the protein catalyst, the process of its folding, and its conformational stability [14]; these structural changes may disturb the reaction flux if they affect the active site [14–18]. Due to these opposing effects, there may be an optimal cytosolic density where cellular efficiency and hence fitness are maximal.

The effect of crowding depends on the size of the catalyst and its substrate: in the presence of other volume-excluding cosolutes, the larger the size of a solute, the stronger the reduction of its diffusion [19] and the perturbation of its free energy [20,21]. Hence, when the size distribution of the molecules changes, the cell needs to adjust its cytosolic density in order to optimize its physiological efficiency.

In a pioneering theoretical study, Vazquez [22] considered how the objective flux of a metabolic network is affected by crowding, assuming that the enzymes are also the crowders in their own right. The corresponding model accounts for the slowdown of diffusion due to crowding, but ignores the growth-rate dependent role of the ribosomal sector and the corresponding changes in the molecular size distribution. This study found that there exists an optimal cytosolic density, which maximizes reaction fluxes and depends on details of the network, such as the ratio between diffusion limited and transition-state limited reactions. Using an alternative modeling approach, Dill et al. also arrive at a similar conclusion [23].

While these studies demonstrated the existence of a flux-optimizing cytosolic density, natural selection maximizes fitness, not metabolic reaction fluxes. For non-interacting cells in a uniform environment, fitness is closely related to the growth rate [24]. The cellular physiology at maximal growth rate—and hence maximal fitness—can be described mathematically through growth balance analysis (GBA) [9].This modeling framework simulates the balanced growth of a self-replicating bacterial cell while accounting for the major physicochemical constraints on cellular growth: mass balance of metabolism and protein production, non-linear reaction kinetics that depend on the concentrations of catalysts and their substrates, and the effects of molecular crowding. Vazquez [22] assumed that all crowding effects can be quantified by classifying reactions into two types: (1) those in the saturation regime, with [*S*] ≫ *K*_M_, and (2) those in the diffusion limited regime, with [*S*] ≪ *K*_M_. This approach constrains the modeled substrate concentrations of each reaction to be either much smaller or much larger than the corresponding *K*_M_, which is incompatible with a realistic modeling of intracellular metabolite concentrations and their effect on molecular crowding [25].

Standard GBA assumes a given, hard limit on dry mass density [9,10] or on total protein concentration [8]. Here, to assess the effects of molecular crowding on fitness, we apply a generalization of GBA that instead describes the kinetics of metabolic reactions and protein translation through crowding-adjusted Michaelis-Menten kinetics [21,26]. We maximize the balanced growth rate while varying the concentrations of transporters, catalytic proteins, and metabolites; these molecular species also form the volume excluding co-solutes in the background of each reaction, affecting diffusion, free energies, and hence reaction kinetics through molecular crowding. Consistent with experimental observations, we find that the cytosolic density of optimal growth strategies depends on the cellular growth rate.

## Results

### Crowding-adjusted reaction kinetics

The effects of molecular crowding on biochemical reaction kinetics are due to volume exclusion effects. Thus, the relevant parameter for their quantification is not the total mass density (dry mass per volume) but the total volume occupancy *ρ*, i.e., the fraction of cytosolic volume occupied by dry mass. However, as molecular mass and volume are approximately proportional [27], we treat density and occupancy as interchangeable, subject to a scaling coefficient that quantifies the mass/volume ratio of the cytosolic dry mass. Depending on the external conditions, the volume occupancy of biopolymers in the cytosol of *E. coli* lies within a range 0.16 - 0.36 [28], providing a lower limit of the total volume occupancy.

To model biochemical reaction kinetics as a function of molecular crowding, we use a modified description of irreversible Michaelis-Menten kinetics [21,26] (see Methods for details),

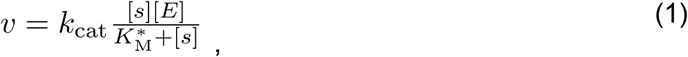

where [*S*] and [*E*] are the concentrations of the substrate and catalyst, respectively; *k*_cat_ is the catalytic rate constant (turnover number); and *K*_M_* is the crowding-adjusted Michaelis parameter [21,26]. In a typical catalytic reaction, a substrate molecule *S* encounters a catalyst molecule *E* to form a catalyst-substrate complex *ES*, which either proceeds forward to convert the substrate into the product, or reverts back to release the substrate. To derive *K*_M_*, we assume that a catalytic reaction can be divided into two subsequent, independent steps that both depend on the cytosolic volume occupancy: (1) *S* and *E* diffuse until their encounter (Fig. 1B) and thereafter (2) they bind and unbind reversibly until the reaction proceeds forward and product *P* is released (Fig. 1A). In step (2), we further assume that *S* and *E* will stay in close proximity and do not diffuse away from each other. These approximations simplify the model derivation and make the reaction times of the two steps additive [21]. Combining the rate laws of the two steps provides an estimate for the effective Michaelis parameter *K*_M_* [21,26]:

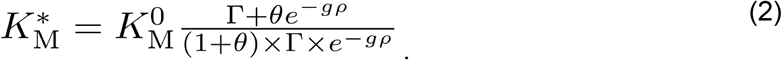

**Figure 1.**
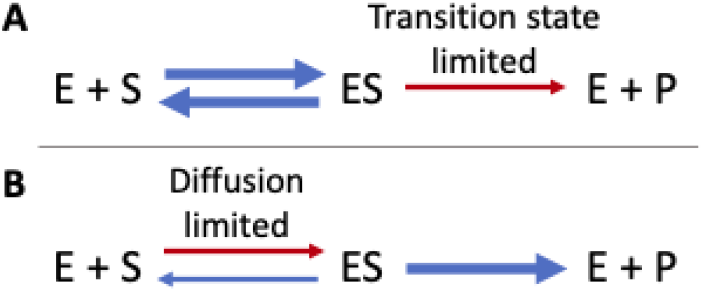
Limiting cases of crowding effects on kinetics. **(A)** Transition state limit, valid if the conversion of enzyme (E) plus substrate (S) to the complex ES is much faster than the conversion of ES to product (P). **(B)** Diffusion limit, where the conversion of ES to P is much faster than the formation of ES.

Here, *K*_M_^0^ is the Michaelis parameter in the low-crowding limit; *Γ* is the correction term for the shift in Gibbs free energy Δ*G* due to molecular crowding; the exponential term exp(-*gρ*) accounts for the slow-down of diffusion, where the scaling factor *g* can be estimated from the size of the substrate, i.e., *g* = *g*(*r*_S_) for a spherical substrate with radius *r*_S_; and *θ* is the relative weight between the rate laws of steps (1) and (2), or in other words the relative ratio between time spent on step (1) and on step (2) at low cytosolic occupancy. Transition state limitation dominates when *θ* is large (θ → ∞), and diffusion limitation dominates when *θ* is small (θ → 0). The best available estimate for *θ* is 2.3, obtained for the ERK MAP kinase phosphorylation reaction [26]; we employ this value in our simulations for metabolic and ribosomal reactions. Small metabolites, however, may diffuse more readily, and so metabolic reactions may have a stronger bias towards being transition state-dominated than ribosomal reactions. Therefore, we also employ an alternative model, in which the metabolic reaction has a *θ* twice as large as that of ribosomal reactions (*θ*=4.6 for metabolic and *θ*=2.3 for ribosomal reactions), and examine if this alternative assumption affects our conclusions.

### Protein translation favors lower occupancy compared to metabolic pathways

To understand the effect of crowding on catalytic reactions, we first consider a simple, linear biochemical pathway model consisting of *N*=20 consecutive enzyme-catalyzed reactions at steady state, i.e., with identical flux through each reaction (Fig. 2A). The kinetics of each step in the pathway depend not only on the concentrations of the catalyzing enzyme and the substrate of the reaction, but—through the crowding effects on *K*_M_*—also on the concentrations of the molecules involved in the remaining *N*-1 reactions. We identified the combination of enzyme and metabolite concentrations that maximizes the pathway output per dry mass, calculated from crowding-adjusted kinetics; adding these concentrations, weighted by the respective molecular volumes, resulted in an estimate of the optimal cytosolic occupancy *ρ*_opt_ for this system. To restrict the number of model parameters, we assumed identical enzyme and substrate sizes as well as identical crowding-adjusted kinetics (Eq. (1)) for all reactions in the pathway. We fixed the physicochemical parameter *K*_M_^0^=130μM, which is the median value for metabolic enzymes and their substrates [29] and is also close to the value estimated for the binding of ternary complexes to the ribosome [30], 120μM. The second physicochemical parameter in this model, *k*_cat_, appears merely as a scaling factor of the pathway flux, and for simplicity we set *k*_cat_=1s^-1^ without loss of generality.

**Figure 2.**
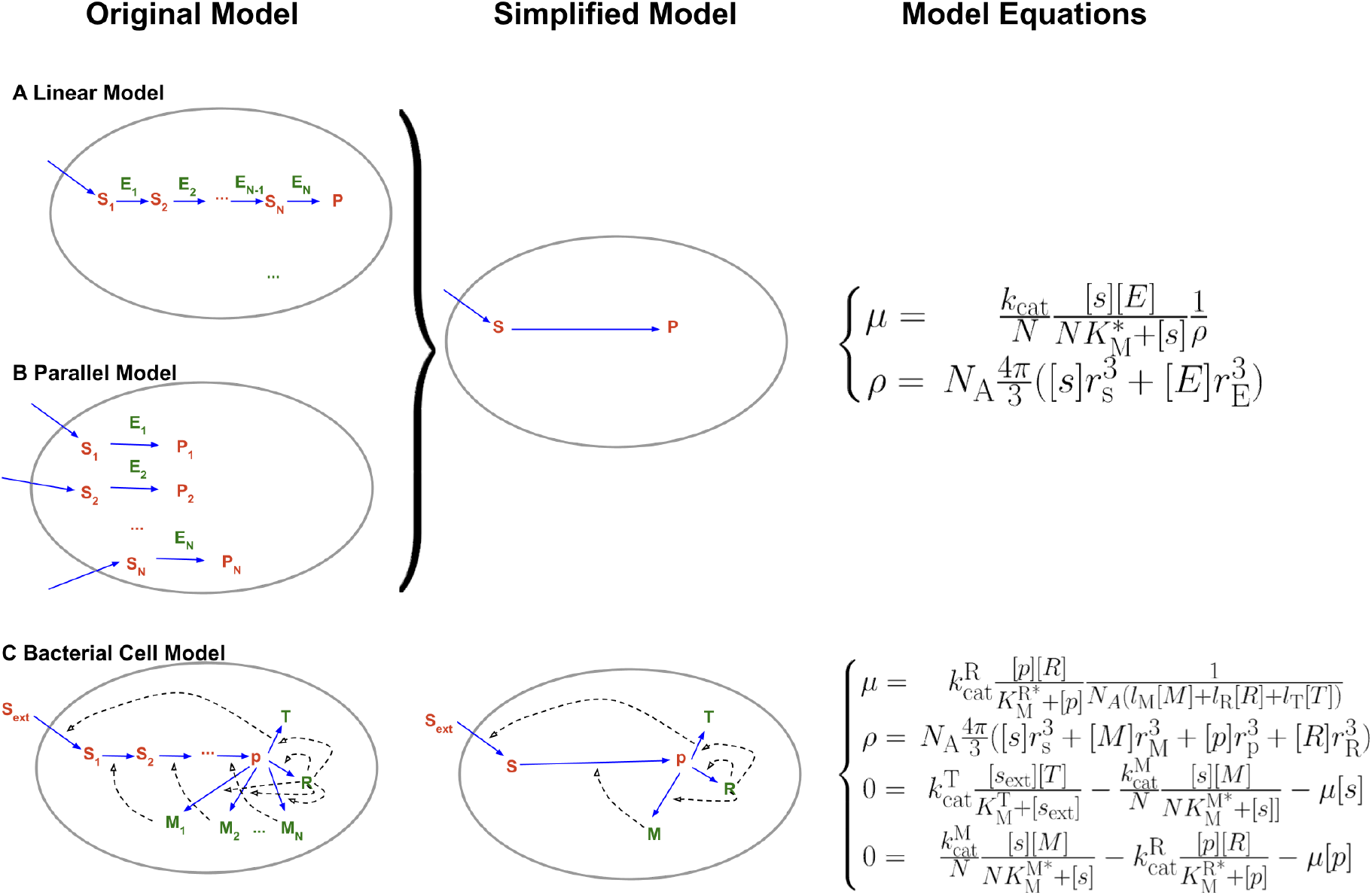
Illustration of the models. **(A) Reaction system in an *N*-steps linear pathway,** representing a metabolic system. The initial substrate, *s*_1_, is replenished by a transport process not included in the model, and is converted by the pathway into the product *p*. **(B) Reaction system of *N* parallel reactions**, representing ribosomes presenting *N* distinct anticodons; *s*_1_,…, *s*_N_ are the corresponding ternary complexes, *p*_1_, …, *p*_N_ are the extending amino acid chains. Assuming that (i) all reactions follow identical kinetics, (ii) all substrate concentrations *s_i_* are identical, and (iii) all enzyme concentrations are identical, the fluxes of the models in (A) and (B) are both mathematically identical to the simplified model to the right of the two panels, defined in Eq. (S7) and Eq. (S8), with re-scaled concentrations [*s*]=[*s*_1_]+[*s*_2_]+…+[*s*_N_] and [*E*]=[*E*_1_]+[*E*_2_]+…+[*E*_N_]. **(C) GBA model simulating the balanced growth of a bacterial cell.** Transporter *T* imports nutrient *s*_1_, which is converted to the precursor for protein production *p* by a metabolic pathway with *N* consecutive enzymes, *M*_1_, …, *M*_N_, via intermediate substrates *s*_2_, …, *s_N_*. The ribosome *R* synthesizes the *N*+2 proteins (*T*, *R*, *M*_1_, …, *M*_N_) from *p*. Assuming identical concentrations of metabolic enzymes, [*M*_1_]=…[*M_N_*]=:[*M*], and of metabolites, [*s*_1_]=…[*s*_N_]=:[*s*], the solution space of this model cell is spanned by the five concentrations [*s*],[*p*],[*T*],[*M*],[*R*] and satisfies Eq. (S9), Eq. (S10), and Eq. (S11). Here *l*_T_, *l*_M_, and *l*_R_ are the number of precursor molecules required to synthesize a transporter protein, a metabolic enzyme, and a ribosome in the model, with values 300, 300, and 7.459, respectively.

We varied the sizes of the catalyst and its substrate, which we assumed to be both spherical. We considered (i) a metabolic pathway, where substrates have sizes typical for metabolites (*r*=0.34nm) and catalyst have sizes typical for globular proteins (*r*=2.4nm); and (ii) a ribosomal system with sizes resembling those of tRNAs (*r*=2.4nm) and ribosomes (*r*=13nm). We also assumed the catalyst-substrate complex to be spherical, with a volume equal to that of the catalyst plus the substrate. Note that while cells do not contain pathways of consecutive ribosomes, the model for the 20-step linear pathway depicted in Fig. 2A is mathematically identical to the model for 20 parallel reactions shown in Fig. 2B if all metabolite concentrations are assumed to be identical; this latter model can be interpreted as a molecular snapshot showing 20 types of ribosomes, each presenting an anticodon for a different tRNA species.

Fig. 3A plots the reaction fluxes. In the metabolic system (blue), the total reaction flux increases monotonically with the cytosolic occupancy within the range explored; note that the term that describes the approximate effects of diffusion, exp(-gρ), breaks down close to ρ=1 [12,26]. In contrast, in the ribosomal system (red), the total reaction flux reaches a maximum at *ρ*=0.21, beyond which potential flux increases due to more and more highly saturated ribosomes are drowned out by increasingly difficult diffusion. Fig. 3B plots the flux per biomass investment, a proxy for growth rate, defined as the reaction fluxes divided by cytosolic occupancy. Here, both systems show a clear optimum, with an optimal occupancy of *ρ*_opt_=0.30 for the metabolic system and *ρ*_opt_=0.12 for the ribosomal system. In the light of natural selection on the cellular growth rate, optimal growth – which occurs at a different occupancy than maximal flux – was likely more relevant.

**Figure 3.**
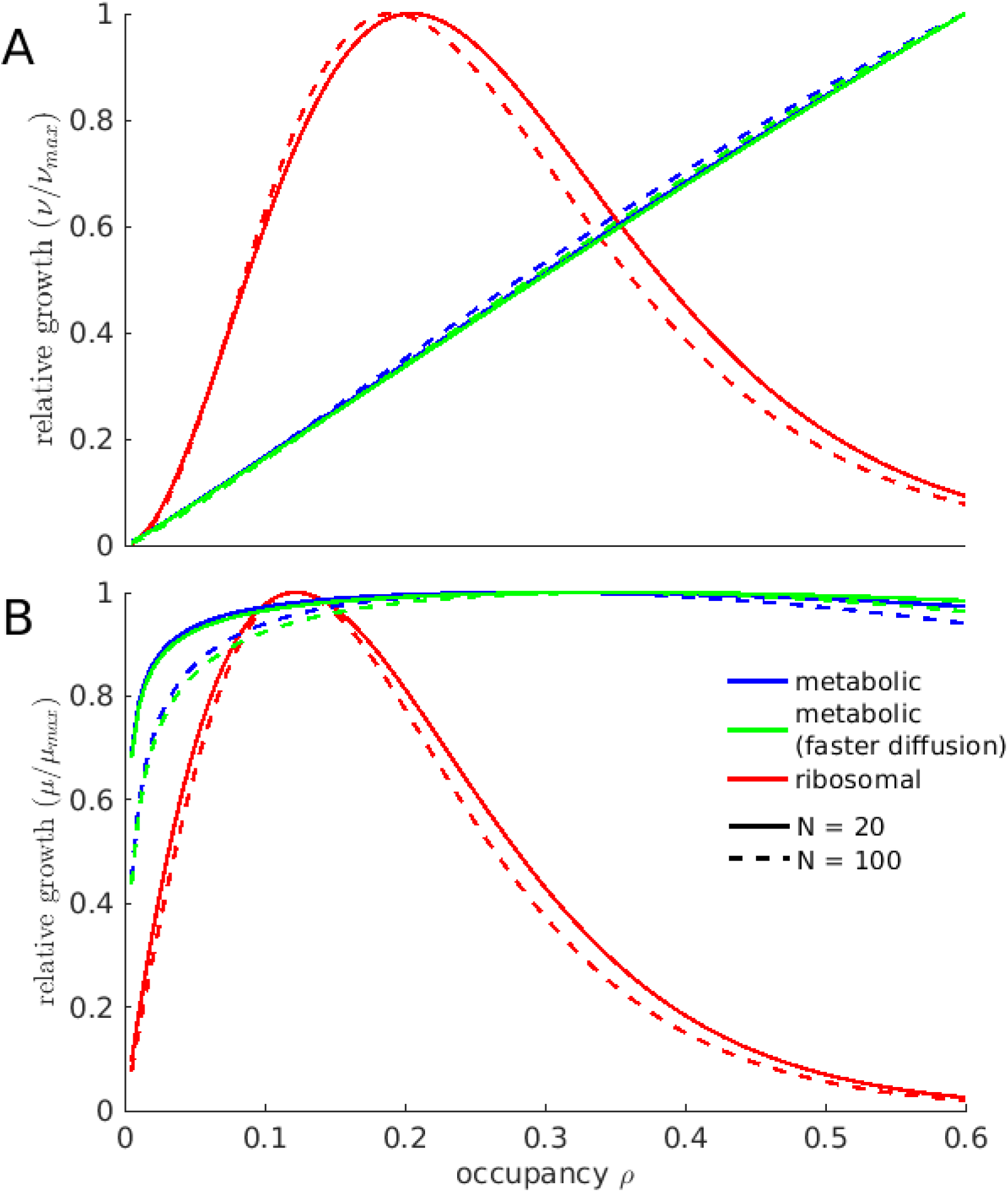
The cytosolic occupancy that facilitates maximal biochemical reaction fluxes is lower for ribosomal than for metabolic systems. (A) reaction fluxes and (B) growth rate (flux per unit dry mass) of the linear model. Blue and green lines represent metabolic systems, with small catalysts and substrates, where blue is for *θ*=2.3 and green is for *θ*=4.6 (stronger bias towards transition state limitation due to fast diffusion). Red lines represent ribosomal systems with much larger molecules. Solid lines are for systems of *N*=20 consecutive (metabolic, blue) or parallel (ribosomal, red) reactions, dashed lines are for larger systems of *N*=100 reactions.

The effect of the occupancy on the pathway output is substantial in the ribosomal system, where a 50% drop in occupancy from *ρ*_opt_ decreases pathway flux per dry mass by 21%, while it is small for the metabolic system, where a 50% drop in occupancy incurs a flux decrease per dry mass of only 1.3%. Increasing the number of steps in the pathway from 20 to 100 has almost no effect on the optimal occupancies *ρ*_opt_ for both systems; however, the reduction of flux per dry mass when *ρ* deviates from *ρ*_opt_ is substantially increased for longer pathways or more parallel reactions (Fig. 3, dashed lines). Assuming that metabolic reactions are more biased towards being transition state limited (green curves in Fig. 3) has only very small effects. Moreover, the approach to crowding employed in Vazquez [22] also leads to conclusions that are qualitatively similar to Fig. 3 (Suppl. Fig. S1). We conclude that the model predictions regarding the differences between metabolic and ribosomal systems are robust and do not depend on model details.

We conclude that the results from the pathway model are consistent with the experimentally observed trend that the cytosol of fast-growing bacteria, which is dominated by the larger molecules of the ribosomal sector, favors a lower occupancy than the cytosol of a slowly growing cell dominated by smaller metabolites and enzymes. Moreover, our results suggest that compared to cells dominated by the ribosomal sector, cells dominated by the smaller molecules of the metabolic sector may suffer a smaller reduction in biochemical efficiency when the cytosol shifts away from *ρ*_opt_.

### Optimal occupancy for a self-replicating cell at balanced growth

Can the single-pathway results be generalized to more realistic models of cellular growth, which combine metabolic and ribosomal activities, and where we can directly assess the effect of concentration changes on cellular growth rates? How does the cytosolic occupancy optimal for growth, *ρ*_opt_, change when the number of active metabolic reactions changes, e.g., when switching from a minimal medium, where all biomass components have to be synthesized from a single carbon source through multi-step biochemical pathways, to a rich medium that provides many cellular building blocks through simple transport processes?

To answer these questions, we considered a schematic GBA model of a bacterial cell [9], the cytosol of which comprises interdependent ribosomal and metabolic sectors (**Fig. 2C**). In this model cell, a nutrient *s*_ext_ is imported into the cell by transport protein *T*; the nutrient is then converted into precursor *p* by an *N*-steps metabolic pathway; finally, the ribosome *R* uses *p* to synthesize all catalytic proteins, including *T*, *R*, and the metabolic enzymes *M*_1_ to *M_N_*. For simplicity, all molecules are again assumed to be spherical in shape. All macromolecules, except for the transporter protein *T*, are located in the cytosol and are thus crowders in their own right; *T* is assumed to be fully integrated into the membrane and does not contribute to crowding. As before, the molecules of the metabolic sector are small (metabolites *s_i_*, with *r*_s_=0.34nm and metabolic enzymes *M_i_* with *r*_M_=2.4nm), while the constituents of the ribosomal sector are much larger (precursor *p* with *r*_p_=2.4nm and ribosome *R* with *r*_R_=13nm).

To estimate the appropriate number of active enzymatic reactions in the metabolic sector, *N*, we used flux balance analysis constrained by enzyme concentration [31]; simulations were performed with an improved implementation parameterized for a genome-scale model of *Escherichia coli* metabolism [32]. We found 259 active metabolic enzymes for growth in a minimal medium with glucose as the sole carbon source; 206 active enzymes for the same medium supplemented with amino acids; and 174 active enzymes for growth in a rich medium (Methods).

To facilitate the numerical determination of the state with maximal growth rate, we approximate the metabolic pathway through a single, lumped reaction with catalyst *M* and substrate *s*, scaling kinetic parameters and molecular masses to account for the pathway length *N*; this approximation neglects the dilution of intermediate metabolite concentrations through volume growth (Methods). The solution space of the model with *N* metabolic enzyme-catalyzed reactions is spanned by 5 concentration variables (**Fig. 2C**), describing the transporter protein with concentration [*T*], the cytosolic substrates with identical concentrations [s_1_]=…=[s_N_]=:[*s*], the protein precursor (“ternary complex”) with concentration [*p*], the metabolic enzymes with identical concentrations [*M*_1_]=…=[*M*_N_]=:[*M*], and the ribosome with concentration [*R*].

The metabolic and ribosomal reactions are described by crowding-adjusted irreversible Michaelis Menten kinetics (Eq. (1)), whereas the transporter reaction is described by conventional irreversible Michaelis-Menten kinetics with constant *K*_M_^⊤^. We investigated how the cytosolic occupancy that allows the fastest growth varies with two parameters, *N* and [*s*_ext_]. *N* is the number of enzyme species in the metabolic pathway, and is inversely related to the richness of the nutrient composition and entering the model through scaling the lumped metabolic reaction. [*s*_ext_] is the concentration of the external nutrient, which parameterizes the degree to which the available nutrients are limiting growth. As above, we challenged the assumptions of our model by performing a second set of simulations, allowing for a stronger bias of metabolic reactions towards transition state limitation (*θ*=4.6) compared to ribosomal reaction (*θ*=2.3).

### Optimal occupancy is lower at faster growth due to higher protein translation demands

In the whole-cell model, the optimal occupancy *ρ*_opt_ generally falls between 0.1 and 0.3 (Fig. 4). *ρ*_opt_ increases when the number of simultaneously active metabolic reactions *N* increases or when the external nutrient concentration *s*_ext_ (and consequently the growth rate) decreases (Fig. 5). These effects are due to shifts in the relative dry mass fractions of the metabolic sector (metabolic enzyme *M* and the substrate *s*; favoring higher occupancy) and the ribosomal sector (ribosome *R* and the precursor *p;* favoring a relatively lower occupancy). Using different *θ* values for the metabolic and ribosomal systems does not change the qualitative trends of the model (Fig. S2). At increasing *N* and constant *s*_ext_, the fraction of the cytosolic volume occupied by the metabolic sector expands at the expense of the ribosomal sector (Fig. S4A), and the saturation of the ribosome with its substrate drops, whereas the saturation of the metabolic enzymes with their substrates varies within a small range (Fig. S4B). At constant *N*=250 and improving nutrient conditions (increasing *s*_ext_), the fraction of the cytosol occupied by the metabolic sector also expands slightly at the expense of the ribosomal sector (Fig. S5A), while the saturation of both metabolic enzymes and ribosomes increases (Fig. S5B). Note that while the cytosolic concentration of ribosomes decreases with increasing growth rate (Fig. S6B), the ribosomal proteome fraction increases (Fig. S6C), as the model cell re-allocates resources from metabolic enzymes and transporters into ribosomes, mirroring experimental observations [1,33].

**Figure 4.**
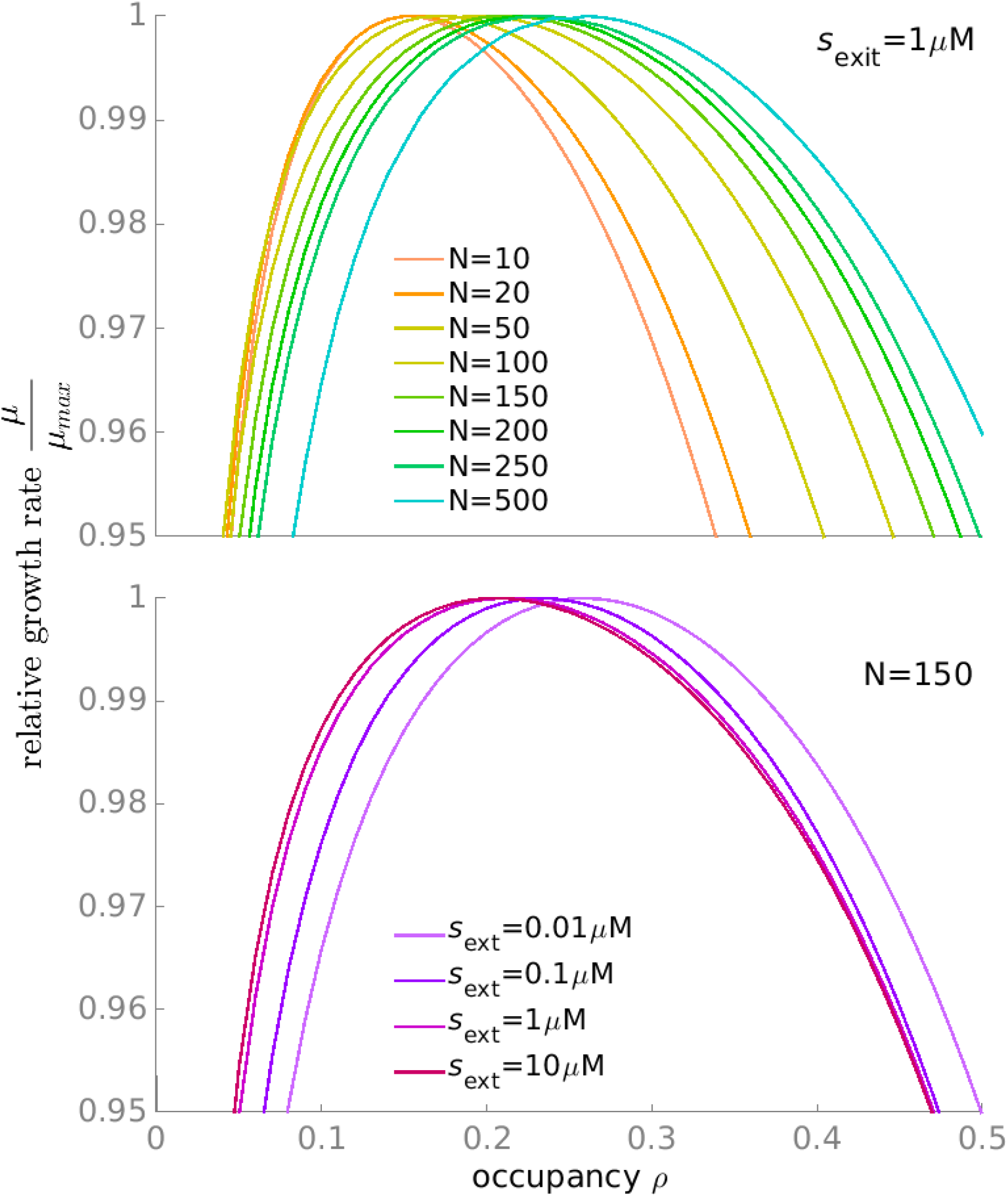
The growth rate *μ* of the crowding-adjusted whole-cell model constrained at different cytosolic occupancies *ρ*. The optimal cytosolic occupancy *ρ*_opt_ **(A)** increases with *N* (the number of enzymes in the metabolic pathway) and **(B)** decreases with external nutrient concentration [*s*_ext_]. *μ*_max_ is the maximal growth rate for each curve across occupancies.

**Figure 5.**
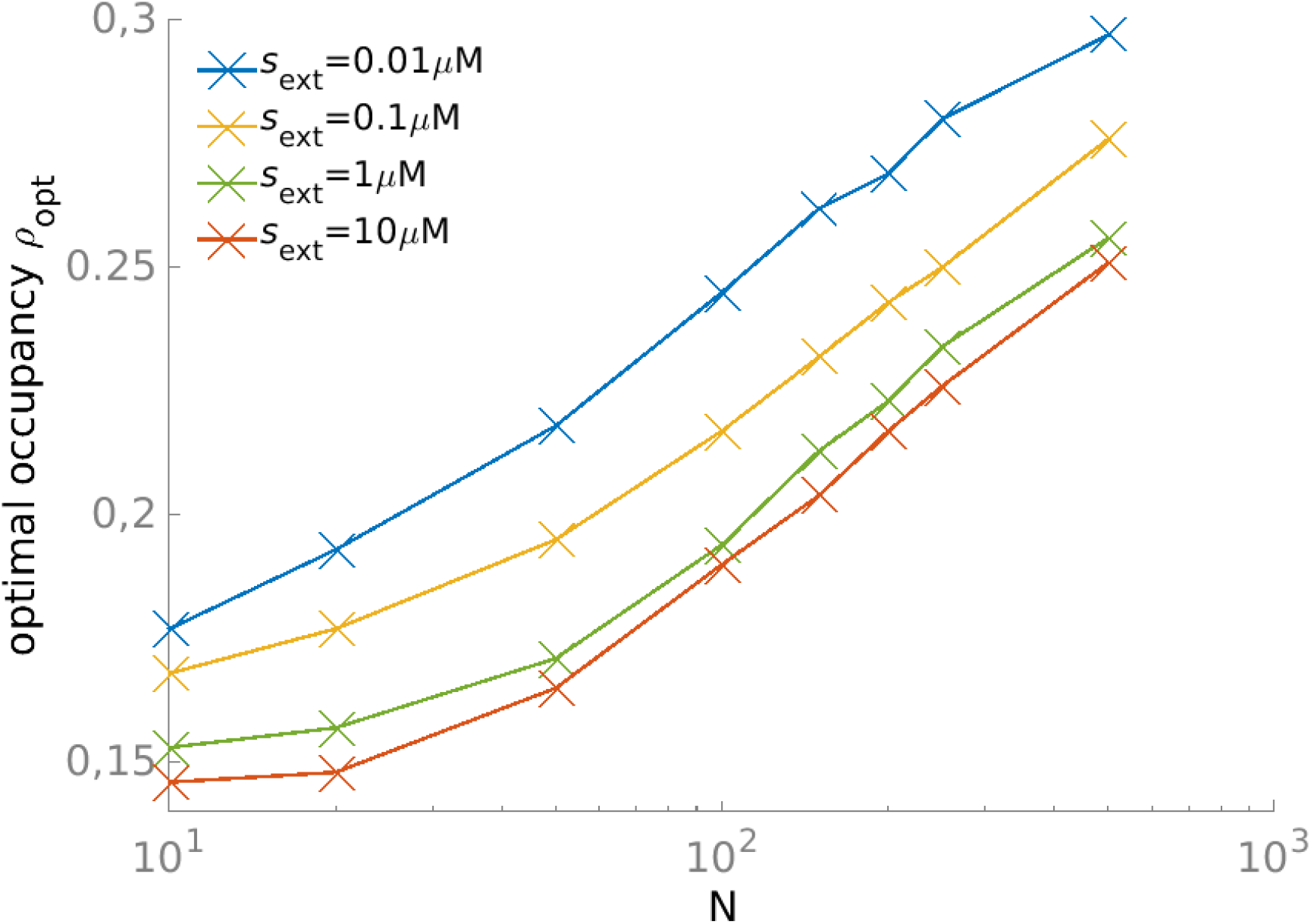
The optimal cytosolic occupancy increases with metabolic pathway length *N* and decreasing external nutrient concentration *s_ext_*.

At optimal occupancy *ρ*_opt_ for different *N* and *s*_ext_, the crowding-adjusted Michaelis parameter *K*_M_* of the ribosomal reaction shows a marked increase with *ρ*_opt_ (Fig. 6; two-sided Spearman rank correlation coefficient *r*=0.995, *P*<10^-15^), consistent with our observations of a strong dependence of the ribosomal flux on *ρ* in the simple pathway model (Fig. 3). In contrast, the *K*_M_* of the metabolic reactions is almost invariant when plotted against *ρ*_opt_ (*r*=0.034, *P*=0.85). These observations support the intuitive notion that the optimal occupancy in the whole-cell model reflects a tradeoff between the saturation of metabolic enzymes, favoring larger occupancies with higher encounter rates, and the inhibition of the ribosomes, favoring lower occupancies with unhindered diffusion of tRNAs. In agreement with our findings for the simple pathway model, the dependence of the growth rate on *ρ* appears to be moderate: e.g., at *N*=150 and *s*_ext_=1μM, a 50% reduction of *ρ* from *ρ*_opt_ reduces *μ* by only 1.1% (Fig. 4).

**Figure 6.**
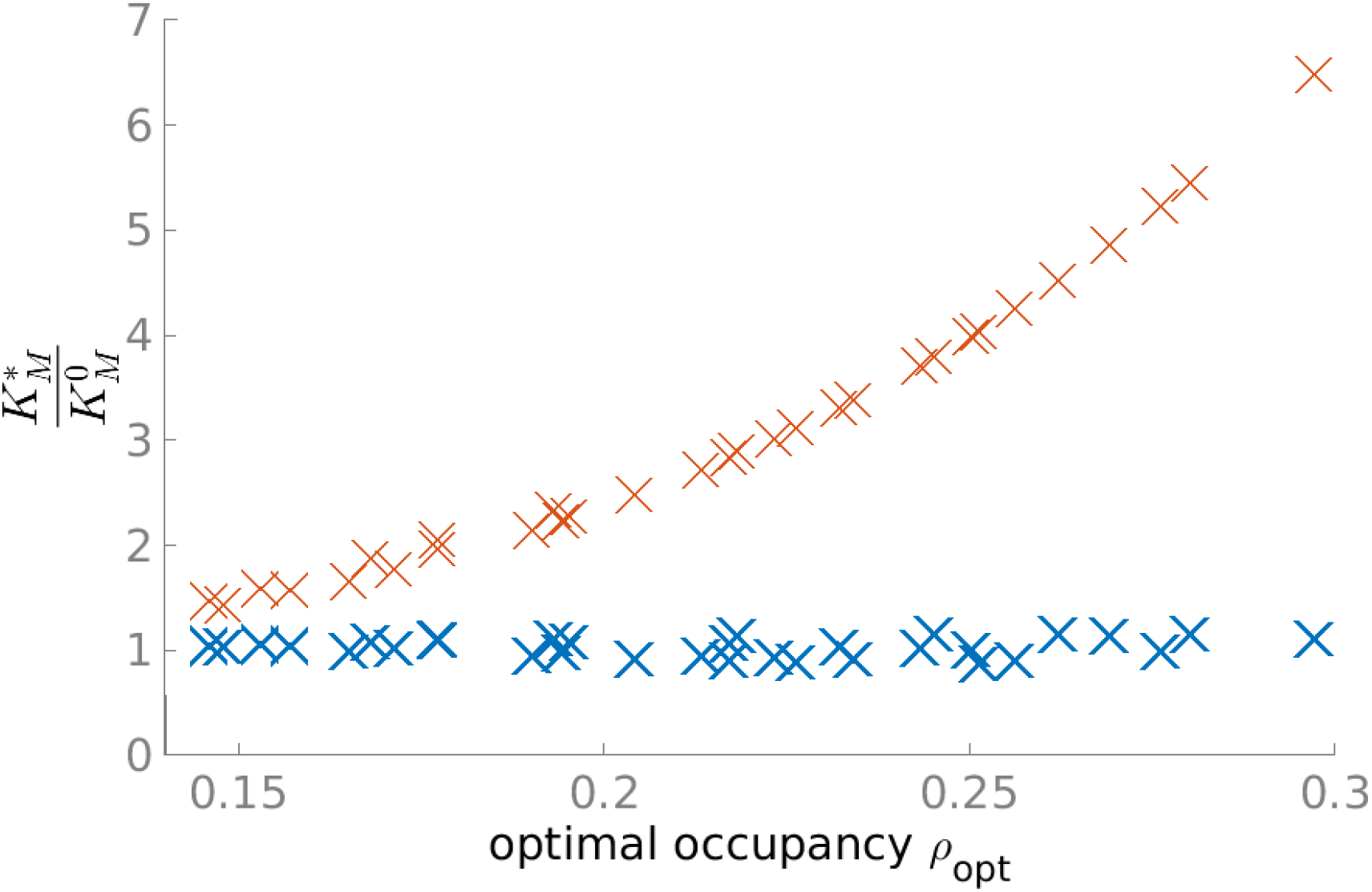
The optimal occupancy strongly influences the effective Michaelis parameter *K*_M_* of the ribosomal (red) but not of the metabolic (blue) reactions in the whole-cell model. Each data point corresponds to a different combination of external nutrient concentration *s*_ext_ and number of active metabolic reactions *N*. While the *K*_M_* of the metabolic reactions does not correlate with *ρ*_opt_ (blue markers; two-sided Spearman’s rank correlation coefficient *r*=0.034, *P*=0.85), the *K*_M_* of the ribosomal reactions correlates with *ρ*_opt_ (red markers; *r*=0.998, *P*<10^-15^). The discrete distribution of points along the *x*-axis reflects the step size used for *ρ* in the simulations.

The slow-down of diffusion and the perturbation of Gibbs free energies have opposing effects on reaction efficiencies. To consider these two effects separately, we define the transition state-perturbation only Michaelis parameter as given by Eq. (2) when setting the diffusion scaling exponent to *g*=0, and we define the diffusion-perturbation only Michaelis parameter as given by Eq. (2) when setting the Gibbs perturbation term to *Γ*=1. We used these hypothetical Michaelis parameters in renewed simulations of the pathway models of the metabolic and ribosomal systems (Fig. 2A,B). In the diffusion-perturbation only model, crowding increases *K*_M_* and reduces the reaction fluxes (Fig. 7, dotted lines); in contrast, crowding in the transition state-perturbation only model decreases *K*_M_* and boosts the fluxes (Fig. 7, dashed lines). At high occupancies (*ρ*≥0.5), the transition state-perturbation effects reach a plateau, whereas the diffusion-perturbation effects continue to increase. Thus, at larger occupancies, the flux-reducing slow-down of diffusion always dominates over the flux-enhancing perturbation of Gibbs free energy when considering their joint effect (Fig. 7, solid lines). In addition, shifting the bias toward stronger transition state perturbation (from solid blue to solid green) delays but does not change the overall trend of *K*_M_* increases with increasing occupancy ρ. Comparison of Fig. 7A and B shows that these trends are largely independent of *N*, the number of reactions in the system. While the trend for the slow-down of diffusion depends on the sizes of the reacting molecules, the trend for the perturbation of Gibbs free energies is largely independent of molecule sizes.

**Figure 7.**
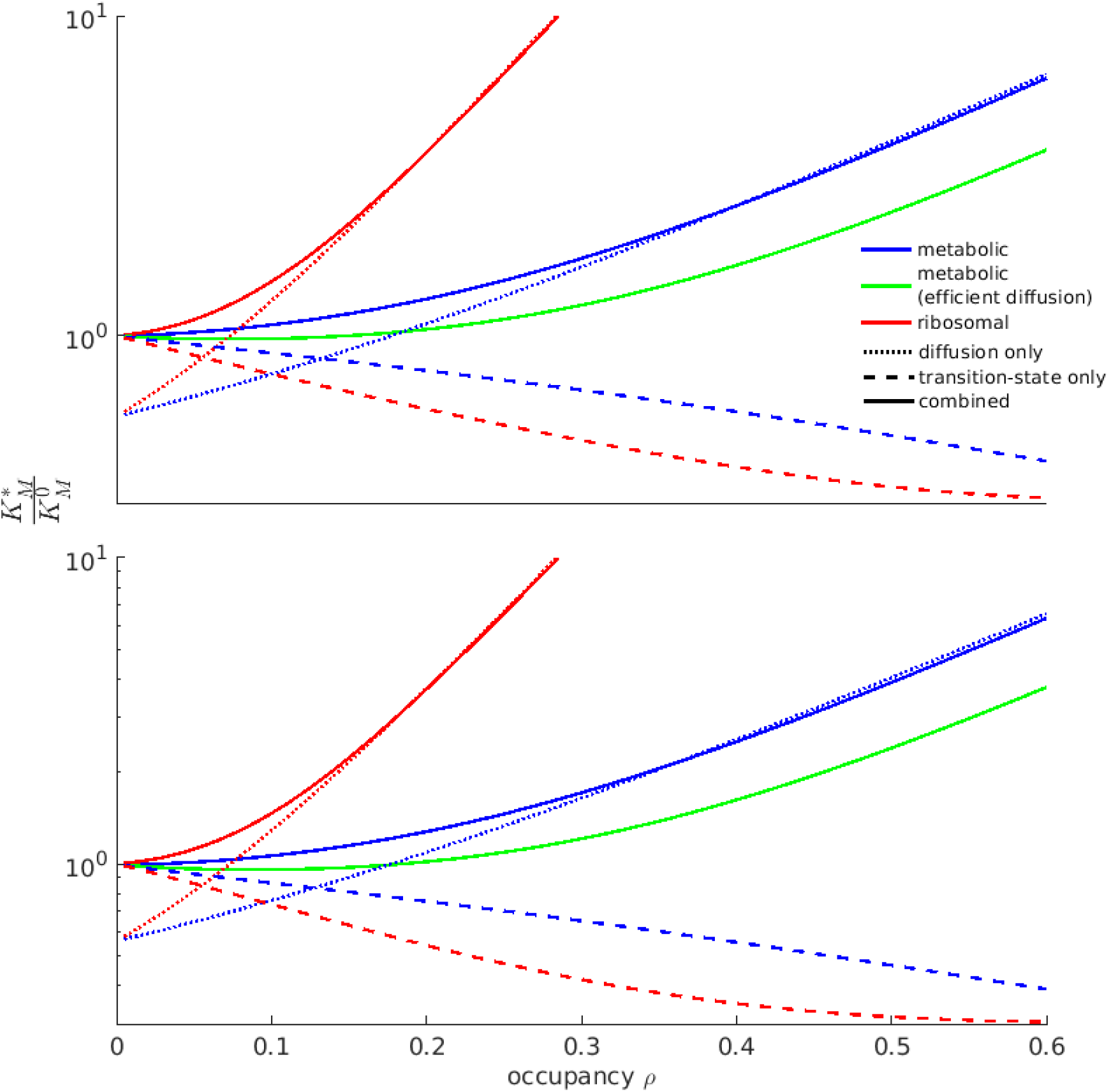
Opposing effects of molecular crowding on the Michaelis parameter *K_M_\** through perturbations of diffusion and of Gibbs free energies. Simulations considered the individual and combined effects of diffusion limitation and transition state limitation for a parallel system of ribosomal reactions (red) and for a linear systems of metabolic reactions (blue: *θ*=2.3; green: combined effect with a stronger bias towards transition state limitation, *θ*=4.6). **(A)** *N*=20 and **(B)** *N*=100.

### Optimal occupancy is quantitatively consistent with the observed E. coli dry mass density

To compare our predictions of optimal occupancy to experimental data from *E. coli*, we first consider the transition from slow to fast growth on minimal media (simulated extracellular nutrient concentration *s*_ext_=0.1μM vs. 1μM, at a constant number of active metabolic reactions *N*=250). Here, the optimal cytosolic volume occupancy *ρ*_opt_ predicted by our whole-cell model decreases from 0.250 down to 0.234 (a 7% drop, Fig. 5).

While no direct experimental observations of the occupancy *ρ* are available, occupancy is expected to scale approximately in proportion to the cytosolic dry mass density *ρ*_DM_ (Lee, 1983). Empirical observations indicate that *ρ*_DM_ does not vary noticeably with *μ* across minimal media, i.e., *ρ*_DM_=0.31g/mL (with 5% measurement error) when *μ*≤0.7h^-1^ (see Fig. 2B of Oldewurtel et al. [6]). As cellular resources are shifted from protein to ribosomal RNA and tRNA with increasing growth rate, and because RNA is denser than protein, the conversion factor between dry mass and volume changes with growth rate. Taking this into account, we find that the experimental data indicates a decrease of *ρ* from 0.221 to 0.208, a 6% reduction that is highly consistent with our estimate (Methods).

Second, we consider the transition from fast growth on minimal media to ultrafast growth in rich media (simulated extracellular nutrient concentration *s*_ext_ of 1μM vs. 10μM, number of active metabolic reactions *N*=250 vs. *N*=150). Here, the predicted optimal cytosolic volume occupancy *ρ*_opt_ decreases by 15%, from 0.234 to 0.204 (Fig. 5). Empirical observation shows that *ρ*_DM_ in *E. coli* decreases from 0.31g/mL to 0.28g/mL as *μ* increases from 0.7h^-1^ to 1.2h^-1^ [6], corresponding to a reduction in occupancy from *ρ*=0.208 to *ρ*=0.184, an 11% decrease. Thus, the predicted change in *ρ*_opt_ is also consistent with the empirically observed reduction in occupancy in the transition to ultrafast growth.

Our simulations also show that even if the cell remains at the state of optimal occupancy, a large reduction in the nutrient level results in only a small decrease of the ribosome’s saturation with its substrate (0.81 to 0.72, Fig. S5B); at the same time, the ribosome’s Michaelis parameter *K*_M_* increases by 51% (Fig. S5C). This finding is consistent with experimental observations that changes in translation rate per ribosome in *E. coli*, a direct consequence of ribosomal substrate saturation, are much smaller than the simultaneous changes in growth rate [34]. At increasing nutrient levels, a higher growth rate *μ* is facilitated by relocating a substantial amount of the ribosome’s synthesis capacity from transporter proteins to metabolic enzymes and additional ribosomes (Fig. S4C).

## Discussion

The linear pathway model shows that reactions with larger catalysts and substrates favour a lower occupancy than reactions with smaller molecules (Fig. 3). This effect explains the observation of lower optimal occupancies for decreasing pathway length *N* in the GBA model cell: the decrease in pathway length simulates the switch from minimal media, requiring on the order of 260 metabolic enzyme species to convert a small number of nutrients to the full range of cellular building blocks, to increasingly richer media, where progressively more biomass components can be taken up directly from the environment, requiring as few as 140 metabolic enzyme species. With decreasing numbers of active metabolic reactions *N*, the ribosomal sector expands at the expense of the metabolic sector, pushing *ρ*_opt_ to lower values (Fig. S4A). Given the minimalistic nature of the whole-cell model, which assumes that all metabolic reactions follow identical kinetics and approximates protein production through a single Michaelis-Menten type reaction, it is striking that the model not only predicts differences across physiological states, but predicts experimentally observed values [6] quantitatively with an error below 12%.

The whole-cell model predicts that the growth rate *μ* decreases only mildly when *ρ* deviates from *ρ*_opt_; for example, *μ* decreases by only 3% when *ρ* increases to twice the optimal occupancy *ρ*_opt_ (Fig. 4; *N*=150, *s*_ext_=1.0μM). This observation is consistent with an experiment that arrested the volume growth of a yeast cell while cytosolic dry mass continued to accumulate, increasing the concentration of a fluorescent protein—a proxy for dry mass density—to roughly twice the wildtype value [35]. These results suggest that the dry mass accumulation rate largely remains constant throughout the volume growth arrest, even when the cytosol density reaches twice the wildtype level.

In the whole-cell model, a 10% deviation of *ρ* from *ρ*_opt_ results in a 0.02% drop in *μ* (Fig. 4); using growth rate as a proxy for fitness, this corresponds to a selection coefficient *s*=2×10^-4^. The effective population size of most bacterial species is on the order of *N*_e_=10^8^ [36], and we thus have *s* ≫ 1/*N_e_* = 10; accordingly, natural selection would be sufficiently strong to explain the difference in average *ρ*_DM_ observed between the two physiological states. Experiments show substantial between-cell variation in each nutritional condition [6]: the observed dry mass densities show coefficients of variation (standard deviation / mean) of around 5%. According to the whole-cell model, the corresponding difference in occupancy corresponds to a selection coefficient of *s* = 10^−4^ » 1/*N* indicating that this large cell-to-cell variation is not selectively neutral but persists despite negative selection.

Questions about the optimal allocation of protein resources can be addressed by maximizing the growth rate (or flux) in computational models of cellular growth with fixed, crowding-unaware kinetic parameters. While such simulations produce meaningful predictions for the relative amounts of the proteins, they cannot limit absolute protein concentrations [37]. To solve this problem, existing models [9,10] implement hard, phenomenological constraints on the total concentration of protein or cellular dry mass, based on experimental observations that found these to be approximately constant across growth conditions [6,33,38,39]. Our results elucidate the biophysical origin of these observations, showing that the cellular dry mass density represents a compromise between the saturation of metabolic enzymes with their substrates and the effects of reduced diffusion on the effective affinity of the ribosome for its much larger substrate, the ternary complex.

## Methods

### Crowding-adjusted Michaelis-Menten kinetics

Macromolecular crowding affects the flux of a metabolic reaction in multiple ways. It can (i) slow down diffusion; (ii) affect the free energy of substrate, catalyst, and the substrate-catalyst complex and thereby change their relative equilibrium ratios; and (iii) disturb the folding of a protein and affect the shape of the active site. In our modelling framework, we followed the derivation proposed in Minton [21] that systematically accounts for the effects of crowding on metabolic fluxes caused by effects (i) and (ii).

In this section, let us consider the metabolic reaction carried out by an enzyme *E* that converts substrate *S* into product *P*, in the presence of other volume-excluding co-solutes that, collectively, constitute the dry mass of the solution. The metabolic reaction is described by the chemical equation following Michaelis-Menten kinetics:

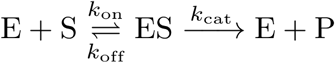

It is described by two parameters: the enzyme-substrate dissociation (or Michaelis) parameter *K*_M_ ~ *k*_off_/*k*_on_, and the catalytic rate constant (or turnover number) *k*_cat_. Note that while *K*_M_ is usually assumed to be invariable and is hence referred to as “Michaelis constant”, we here examine crowding-dependent changes in *K*_M_ und hence refer to it as the “Michaelis parameter”. We assume that all reactions follow effectively irreversible Michaelis Menten kinetics and thus free energy changes are irrelevant to the transition rate *k*_cat_; moreover, we ignore crowding effects on the enzyme structure, and thus *k*_cat_ is not affected by crowding. To derive the effect of crowding on the Michaelis parameter *K*_M_, we divide a catalytic reaction into two independent, consecutive steps: (1) *S* and *E* diffuse until they meet, and then (2) *S* and *E* bind and unbind reversibly until the reaction proceeds forward to make *P.*

#### Step (1): the substrate-catalyst encounter

The encounter rate between *S* and *E* depends on the cytosolic occupancy. Its rate law is similar to that of a diffusion-limited catalytic reaction, in which the *ES* encounter rate is low and their encounter is the rate determining step; ys soon as the *ES* complex is formed, the reaction quickly proceeds, converting *S* into *P* and releasing *E*. At given concentrations of *E* and *S*, the rate of formation of *ES* is thus mostly determined by the rate of encounter, which is proportional to the sum of *E*’s and *S*’s diffusion coefficients. A reduction of the diffusion rate shifts the equilibrium between the concentrations of the individual molecules and the complex *ES*; this can be accounted for in the following way [21]:

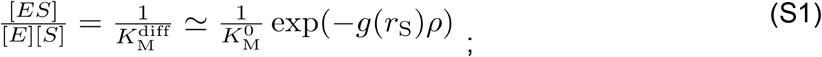

here, *K*_M_^diff^ is the hypothetical diffusion limited Michaelis parameter, while K_M_^0^ is the Michaelis parameter in the low-crowding limit; ρ is the volume occupancy of the volume-excluding co-solutes (dry mass) of the solution (with range 0 < ρ < 1), and g is a function that depends on the shape of E, S, and other volume-excluding co-solutes. Since S is typically much smaller than E, the diffusion coefficient of S in a crowded solution is in general much higher than that of E. Hence, we estimate this scaling term exp(- *g*ρ) solely from the diffusion coefficient of S. Approximating S as a sphere of radius r_S_, we can write it as exp(– *g*(*r*_S_)ρ).

The bacterial cytosol is crowded, slowing down the diffusion of all molecular species. The extent of this slow-down, however, is non-uniform and depends largely on the size of the affected molecule: the larger the molecule, the more it is slowed down. The slow-down of diffusion in the *E. coli* cytosol is summarized by the following empirical scaling law, which was inferred from molecular dynamics simulations [19]:

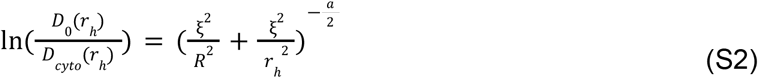

where *r_h_* = 1.3(*r* + 1.4Å) is the hydrodynamic radius of a molecule with radius *r* [40], *i.e.*, its effective radius including the attached water molecules; *D*_0_(*r*_h_) is the diffusion coefficient in the low crowding limit, while *D*_cyto_(*r*_h_) is the diffusion coefficient in the crowded cytosol condition; ξ=0.51nm is the average distance between the surfaces of volume-excluding co-solutes in the cytosol of *E. coli*; *R*=42nm is the radius of the largest common crowders in the cytosol; and *a*=0.53 is an empirical scaling factor [19]. Note that in general, the parameter *ξ* depends on *ρ*; however, as we are only interested in the relationship between reaction fluxes, growth rate, and *ρ* in a small range centered around the native *E. coli* cytosolic density, we can approximate *ξ* by a constant in our analysis. As the cytosolic dry mass density is ~0.3g/mL [6] and the mass-to-volume-ratio of protein is 1.35g/mL [41], the cytosolic volume occupancy *ρ* is approximately 0.22=0.3/1.35. If the reaction rate is proportional to the rate of encounter between *E* and *S*, and this rate of encounter is in turn approximately proportional to the diffusion coefficient of *S*, then Eq. (S2) can be used to calculate the scaling factor *g*(*r*_S_):

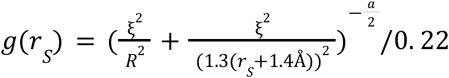

and the rate of this reaction is denoted as *k*_diff_:

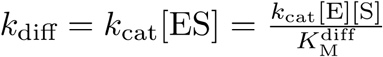

#### Step (2): repeated binding and unbinding of substrate and catalyst

The rate law of this step is related to a transition-state limited catalytic reaction, in which the rate of encounter of E and S is much larger than the rate of conversion of the *ES* complex into the product P. In this case, the complex *ES* exists in near equilibrium with *E* and *S*, and macromolecular crowding affects the reaction rate parameter mainly through shifting this equilibrium. To assess the magnitude of this effect, we consider the reversible reaction

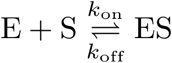

and denote the adjusted equilibrium Michaelis parameter in this hypothetical transition-limited case as *K*_M_^ts^ [21]:

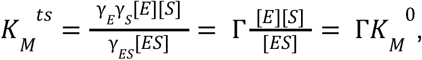

where *γ_i_* denotes the activity coefficient of molecular species *i*, 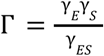, and 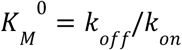 is the enzyme-substrate dissociation parameter in the low-crowding limit. The activity coefficients are defined as

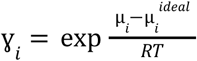

with

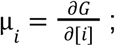

here, G is the Gibbs free energy of the system; [*i*] is the concentration of molecular species *i*; *μ_i_* is the chemical potential of *i*, and 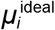 is the chemical potential in an idealized situation, i.e., without intermolecular interactions and in the absence of other volume-excluding co-solutes. In other words, the equilibrium of the reaction will be identical to the dissociation parameter *K*_M_^0^ if the system is ideal. The *Γ*term, therefore, accounts for the deviation of the Gibbs free energy from the idealized situation.

The *Ɣ_i_* of each molecular species *i* can be written as an expansion in terms of the concentrations of all molecular species [42]:

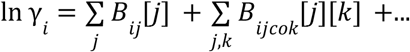

The coefficients *B_ij_* (*B_ijk_*, …) reflect the interaction between 2 (3, …) molecular species. For example, the coefficient *B_ij_* is given as [43]:

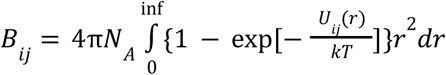

where *N*_A_ is Avogadro’s number, *r* is the distance between the center of mass of molecular species *i* and *j*, and *U_ij_*(*r*) is the potential of average force acting between the two molecular species. While the interaction potential among multiple molecular species is complex, it has been found that the colligative properties of solutions of globular proteins can be well accounted for over a wide range of concentrations by using a simple hard sphere potential, where two molecules cannot overlap but do not interact otherwise (see Minton [21] for a review):

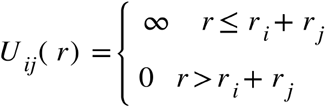

The scaled particle theory applies this rigid sphere potential to calculate the activity coefficient *γ_i_* of molecular species *i* (Boublík, 1974):

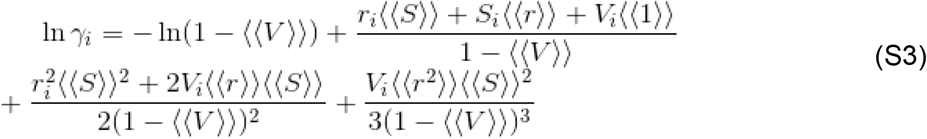

Here, *w*, is the concentration (number density) of molecular species *i*; 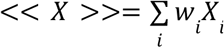 weighted sum of property *X*, with 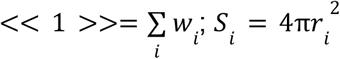, is the surface area and
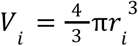 is the volume of molecular species *i*; in addition, 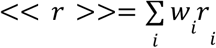 and 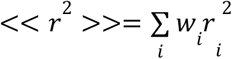. As in the hypothetical transition-state limited case, the reaction rate parameter is proportional to the concentration of the enzyme-substrate complex, and a shift of its equilibrium constant by Γleads to a corresponding shift of the reaction rate, quantified by the rate parameter *k_ts_* [21]:

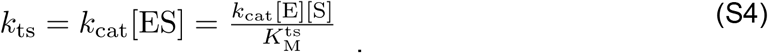

#### Combining step (1) and (2) for the general formulation

We assume that a catalytic reaction is divided into two independent steps that occur in tandem: (1) the encounter of *S* and *E*, whose rate law is proportional to that of a diffusion limited reaction, and thereafter (2) their reversible binding and unbinding until *S* is converted into *P*, whose rate law is proportional to a transition-state limited reaction. The crowding-adjusted reaction rate *k* is obtained by adding the inverse rate parameters (i.e., the reaction times) of the two steps [21,26,44]:

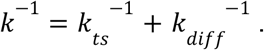

Given Eq. (S1) and (S4), this leads to [26]:

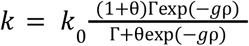

Here, 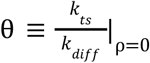 quantifies the relative contributions of step (1) and (2) to the overall reaction, or in other words the ratio between times spent on step (1) and on (2) at low cytosolic occupancy; the reaction tends towards diffusion limited if *θ*→0, or transition-state limited if *θ*→∞. *k*_0_ is the overall rate parameter in the absence of crowding. Remember that [*E_free_*] = [*E_total_*] – [*ES*] = [*E_total_*]/(1 + [*S*]/*K_m_*) is the concentration of free enzymes only, and thus the reaction rate is *k* = *k_cat_*[*ES*] = *k_cat_*[*E_free_*][*S*]/*K_M_*; from now on, we describe the reaction rate as the flux of the reaction, and denote it as *v.* As we assume that the reaction is irreversible and hence crowding does not influence *k*_cat_, to arrive at crowding-adjusted Michaelis-Menten kinetics, we have to scale the Michaelis parameter, which arises from considerations on the equilibrium between the enzyme-substrate complex and its constituents. Thus, 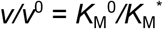, with *K*_M_^0^ the Michaelis parameter in the low crowding limit; then the crowding-adjusted Michaelis parameter is

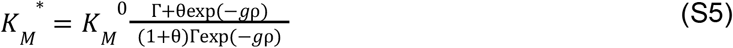

and so the flux can be written as

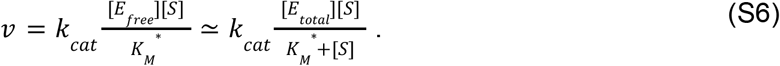

The *θ* of ERK MAP kinase phosphorylation reaction was estimated to be 2.3 [26]; we assume that this value is representative for cellular enzymes and use it for modeled reactions. To understand how the choice of *θ* affects the model behavior, we also ran a model where metabolic reactions have *θ*=4.6.

The concentration of the enzyme-substrate complex, [*ES*], depends on *K*_M_* through [*ES*] = [*E_total_*][*S*]/([*S*] + *K_M_*\*); at the same time, *K*_M_* depends on the concentration of different molecular species, including [*ES*], through Eq. (S3). To find a self-consistent solution for these two quantities, we iterate these two equations until convergence. In each iteration, we first calculate the concentration of the catalyst-substrate complexes of different reactions; we then update their *K*_M_* and proceed to the next iteration. We stop the interaction when all *K*_M_* values changed by less than 0.001% compared to the previous iteration.

### Single-pathway model

We considered a simple model of a linear pathway to investigate how the size of the substrate and catalyst of a metabolic reaction affect the optimal cytosolic occupancy; here, optimality is defined as a maximal pathway flux per unit dry mass, calculated from crowding-adjusted kinetics. The pathway is divided into *N* steps, where *E_n_* is the catalyst of step *n*, which converts its substrate *s_n_* into *s*_*n*+1_, the substrate of the next step (Fig. 2A). We assume that all internal metabolite concentrations are in steady state (i.e., producing and consuming fluxes cancel exactly), and we ignore the dilution of intermediates. Thus, all reaction fluxes have the same value, *v.* We assume that *s*_1_ is replenished by a flux *v*_*s*_1__ = *v*, which is not modeled explicitly.

We assume that the *N* reactions are described by crowding-adjusted Michaelis Menten kinetics with identical *k*_cat_ and *K*_M_^0^. We further assume that the *N* catalyst species and the *N* substrate species are spherical, with radius *r*_E_ (volume *V*_E_) for the catalysts and radius *r*_s_ (volume *V*_s_) for the substrate species. These assumptions simplify the solution space of the model, as in the optimal steady state, all catalysts have equal total concentrations ([*E*_1_]=[*E*_2_]=…=[*E_N_*], and so do all the substrates ([*s_1_*]=[*s*_2_]=…=[*s_N_*]). We define the total substrate concentration 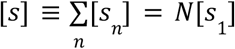 and the total catalyst concentration 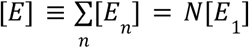. The two variables [*s*] and [*E*] span the solution space of this model. Substrates and catalysts, as well as the substrate-catalyst complexes, are crowders in their own right; they slow down diffusion and perturb Gibbs free energies. The cytosolic volume occupancy of dry mass in the solution, *ρ* (0 ≤ ρ ≤ 1), is

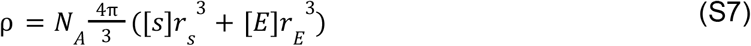

with the Avogadro number *N*_A_. The flux through the pathway per unit volume is

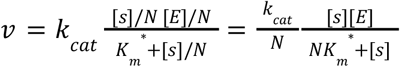

Accordingly, the flux per unit dry mass is

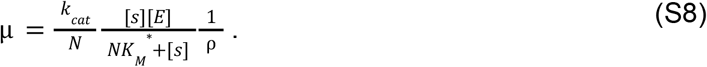

As we ignore crowding effects on the turnover number, *k*_cat_ acts only as a scaling factor, and we thus set *k*_cat_=1 for simplicity. We set the Michaelis parameter at the low crowding limit to *K*_M_^0^=130μM, which is the median *K*_M_ of metabolic enzymes [29], and is also very close to the Michaelis constant of the ribosome estimated from the diffusion limit without molecular crowding [30].

Because we assume identical kinetics of all reactions and ignore the dilution of intermediates through growth, the whole pathway is equivalent to a single reaction with re-scaled kinetics. We still describe it as an *N*-steps pathway, as this more faithfully reflects the situation in the real cell, allowing us to use realistic parameter values. Moreover, while mathematically, *N* represents a scaling factor, it has an intuitive biological interpretation.

The same equations can be used to describe a system of *N* parallel enzymatic reactions with identical fluxes, with only an additional multiplication by *N* in Eq. (S8), *μ*_parallel_=*μ* x *N* (Fig. 2B). Here, catalyst *E_n_* of reaction *n* converts substrate *s_n_* into the end product, and the consumption of *s_n_* is compensated by a flux *v* that supplies *s_n_* at an equal rate.

We consider two systems of substrate and catalyst sizes: metabolic and ribosomal. In the metabolic system, we use *r*_s_=0.34nm for metabolites (the approximate radius of the amino acid alanine [45]) and *r*_E_=2.4nm for globular proteins (an “average” protein in *E. coli* has a mass 40kDa [3], while a globular protein with mass 50kDa has a radius *r*=2.4nm [46]; in an alternative estimation, the radius of a typical globular protein is approximately *r*=2.5nm [47]). In the ribosomal system, we use *r*_s_=2.4nm for the ternary complexes (the gyroscopic radius of tRNA is estimated to range from 2.33nm to 2.46nm based on Eq. (7) of Hyeon et al. [48]). We use a radius of *r*_E_=13nm for the ribosome, as the diameter of a ribosome is reported to be 20nm - 30nm [47,49,50]. We assume that the catalyst-substrate complex is also spherical, with a volume equal to the sum of the substrate’s volume and the catalyst’s volume. For both metabolic and ribosomal systems, we used crowding-adjusted Michaelis-Menten kinetics with *θ*=2.3; to explore the effect of assuming the same *θ* value for both systems, we alternatively examined a model where we set *θ* to 4.6 for the metabolic system, as the metabolic system may have a higher diffusion efficiency than the ribosomal system.

For a fixed value of the total occupancy *ρ*, we calculated the specific flux *μ* using MATLAB, while (i) varying the occupancy *ρ* in steps of 0.01 from 0.01 to 0.8, with additional, finer-grained steps of 0.001 from 0.100 to 0.360, and (ii) varying the ratio of the volume occupied by the substrates *s*, from 0.1% to 97.7%, with an increase by a factor of 1.0023 at each step.

### Single-pathway model: the Vazquez approach

In Vazquez [22], reactions are classified into two contrasting types: (1) those in the saturation limit, with [*S*] *≫ K*_M_, and (2) those in the diffusion limit, with [*S*] ≪ *K*_M_. The rate of reactions in the diffusion limit is modified by an exponential term, exp(- 5.8ρ), where 0≤*ρ*≤1 is the cytosolic occupancy. In addition, crowding increases the contact with enzymes and substrates, and so the reactions are sped up by the term 1/(1 – ρ). Overall, the reaction rate of a reaction in saturation is

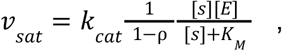

while the reaction rate in the diffusion limit has an extra exponential term,

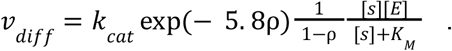

We simulated the metabolic and ribosomal systems based on these two equations, setting *k*_cat_=1 and assuming the metabolic system to be in saturation and the ribosomal system to be in diffusion limit (Fig. S1).

### Model cell with a metabolic and a ribosomal pathway

To more faithfully represent a complete cell and to study the tradeoff between metabolic and ribosomal reactions, we also consider a balanced growth model with a metabolic sector and a ribosomal sector (Fig. 2C). As seen from the results for the pathway models, the two types of reactions have very different optimal conditions: the metabolic sector involves smaller catalysts and substrates than the ribosomal sector, and hence has maximal specific fluxes at a higher cytosolic occupancy.

In this model (Fig. 2C), the transporter *T* imports the external substrate *s*_ext_ into the cytosol, where it is now labeled *s*_1_. The transport reaction is described by ordinary, crowding-unaware irreversible Michaelis Menten kinetics, with flux

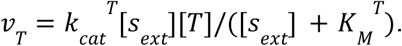

We set *k*_cat_^⊤^=13.7s^-1^ (the median *k*_cat_ of enzyme reactions [29] (Bar-Even et al., 2011)), and *K*_M_^⊤^=1μM (close to the growth limiting glucose concentration of *E. coli* [51]).

The metabolic sector comprises an *N*-steps linear pathway of metabolic reactions, identical to the one studied in the simple pathway model: the enzyme of metabolic reaction *n* (1 ≤ *n* ≤ *N*), denoted as *M_n_*, converts substrate *s_n_* into *s_n_*_+1_; in reaction *n*=*N, M_N_* converts one *s_N_* into one precursor *p* [8]. These *N* reactions follow crowding-adjusted irreversible Michaelis Menten kinetics with identical rate parameters *K*_M_^M0^=130μM and *k*_cat_^M^=13.7s^-1^ (the median *K_M_* and *k*_cat_ of enzyme reactions [29]). As before, all *N* substrates have radius *r_s_*=0.34nm, while all *N* enzymes have radius *r*_M_=2.4nm; as before, we assume that the enyzme-substrate complex is spherical and occupies as much volume as one substrate plus one enzyme molecule. To facilitate the numerical solution of the model, we assume that all metabolite concentrations [*s_n_*] and also all enzyme concentrations [*M_n_*] are identical. This corresponds to the optimal balanced growth solution when the differential dilution of intermediate metabolites is ignored [29]; thus, we treat the dilution of intermediate metabolites only approximately here. In the balanced growth condition, the production rate of each substrate is equal to its consumption rate plus its (approximate) rate of dilution through growth, so that its concentration remains stable. We define the total substrate concentration 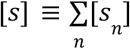 and total enzyme concentration 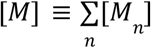. The flux of each metabolic reaction is 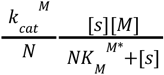.

The ribosomal sector comprises the ribosome (*R*) and the protein precursor (*p*): *R* converts *p* into the *N*+2 types of protein in the model, *N* metabolic enzymes (*M_n_*), the ribosome (*R*), and the transporter (*T*). As before, the radius of *p* is *r_p_*=2.4nm and the radius of *R* is *r_R_*=13nm; their complex is assumed to be spherical and to occupy as much volume as one precursor plus one ribosome molecule. The ribosomal conversion rate is described by crowding-adjusted irreversible Michaelis Menten kinetics with parameters *K*_M_^R0^=120μM [30] and *k*_cat_^R^=22.0s^-1^ [8,52], and the consumption rate of *p* by *R* to make proteins is 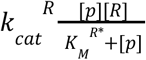. The ribosome converts *l*_⊤_=300 precursors into one transporter, *l*_M_=300 precursors into one metabolic enzyme, and *l*_R_=7459 precursors into one ribosome [8].

Note that the reactions that produce and consume the precursor *p* do not conserve volume. The reason is that to model a realistic cell, we envision the precursor as a charged tRNA, only the amino acid part of which is (i) produced by the metabolic pathway and (ii) integrated into the growing protein. The metabolic pathway only provides the amino acid, while the pool of free tRNAs is not explicitly modeled. For this reason, the size of *p* is substantially larger than the size of the metabolite *s*_N_ consumed in its production. Conversely, the ribosome consumes 300 precursors to produce a single transporter or enzyme, which are both substantially smaller than the combined volume of the 300 precursors. Here, we envision that the tRNA part of *p* is set free again and can be re-charged through *M*_N_. This treatment assumes that the concentration of free tRNA is so low that we can ignore its dilution through growth and its contribution to the cytosolic crowding.

The solution space of this model cell spans five dimensions: [*s*], [*p*], [*T*], [*M*], and [*R*]. The corresponding molecules, along with their complexes, are also the crowders that slow down diffusion and disturb Gibbs free energies. The cytosolic occupancy *ρ*, 0 ≤ ρ ≤ 1, is

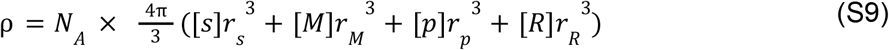

with the Avogadro number *N*_A_. The growth rate *μ* of this model cell can be expressed as the flux through the ribosome reaction divided by the total protein concentration,

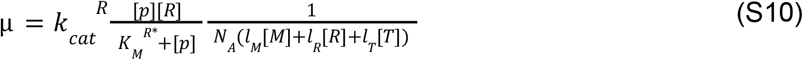

In the balanced growth state, the production of *s* and *p* is offset by their consumption and dilution by growth,

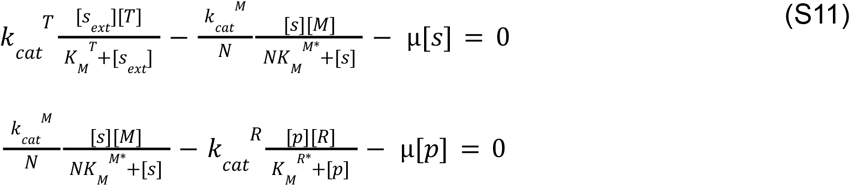

For a given number of enzymes *N* and occupancy *ρ*, we solved this model numerically. As a preliminary step, we used the BARON algorithm [53] implemented in Pyomo [54,55] and assumed normal crowding-unaware irreversible MIchaelis Menten kinetics, maximizing the growth rate *μ* over the space of concentrations ([*s*],[*p*],[*T*],[*M*],[*R*]), subject to the constraints of Eq. (S9) and (S11). In the main model, we assumed *θ*=2.3 for both metabolic and ribosomal reactions. As an alternative, we also examined a model with metabolic reactions biased more toward the transition state limit, setting *θ*=2.3 for ribosomal reactions and *θ*=4.6 for metabolic reactions.

Using the BARON solution as a starting point, we applied the SLSQP algorithm within the function *“minimize”* in SciPy [56], now using crowding-adjusted Michaelis-Menten kinetics. It is not clear if the optimization problem has a unique solution; to increase the probability that the solution is a global maximum, we repeatedly ran the SLSQP algorithm at least 20 times for each (*N,ρ*) combination, and picked the solution with the highest *μ*. For each simulation, at least half of the independent runs supported the same, maximal optimum.

### Estimating the number of enzymes in the metabolic pathway

To obtain a realistic estimate of the number of simultaneously active metabolic reactions in a bacterial cell, we performed flux balance analysis simulations accounting for molecular crowding in terms of a hard limit on the total cellular protein concentration. Simulations were run using “sybil”, an R library for efficient constraint-based analyses [32,57,58]. We used sybilccFBA [59,60], a re-implementation of the MOMENT algorithm with an improved treatment of multifunctional enzymes, which maximizes the biomass production rate while constraining the sum of cytosolic metabolic enzyme concentrations at the experimentally observed level.

We used a sybilccFBA implementation of the iAF1260 stoichiometric model [61], parameterized with turnover numbers for *E. coli* [59,60]. We considered four different nutritional environments. In each case, we counted the number of active metabolic reactions with both substrates and products located in the cytosol, and with a flux >10^-6^ mmol/(gram Dry Weight)/h to filter out numerical noise; we enumerated the enzymes supporting these reactions, filtering out those with proteome fraction lower than a cutoff (10^-9^; for reference, the most abundant enzyme has a density ~10^-3^).

The estimated numbers of active metabolic enzymes and reactions in the different conditions are as follows: (i) 259 active enzymes and 349 active reactions in a minimal medium with glucose as the sole carbon source, corresponding to slow to intermediate growth with a cytosol dominated by the metabolic sector [59,62] (Supplementary Table S1); (ii) 206 active enzymes and 288 active reactions in the same minimal glucose medium supplemented with 20 amino acids, simulating intermediate growth [59,62] (Supplementary Table S2); (iii) 174 active enzymes and 250 active reactions in a rich medium, corresponding to fast growth and a cytosol dominated by the ribosome and its substrates [59,63] (Supplementary Table S3); and (iv) 140 active enzymes and 234 active reactions in an extremely rich medium, where all exchange reactions in the iAF1260 *E. coli* model are allowed to be active.

### Conversion of experimental dry mass density to occupancy

An empirical growth law connects the RNA/protein mass ratio (*r*) of an *E. coli* cell with its growth rate *μ*: *r =* 0. 087 + μ/(4. 5h^-1^) (Eq. (1) of Scott et al. [1]). When *μ* increases from 0 to 0.7h^-1^, *r* increases from 0.087 to 0.243. The average density of protein is 1.35 g/mL [41], while that of the *E. coli* 70S ribosome is (1.637 g/mL), where RNA constitutes 61.87% and the remaining mass fraction is assumed to be proteins [64]. From these relationships, we can obtain the RNA density, resulting in a value of 1.81 g/mL.

If we ignore all molecular species in the dry mass except protein and RNA (which constitute ~75% of the dry mass in an *E. coli* cell [65–67]), we find that the overall density (mass per volume of dry mass) increases from 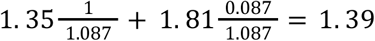 to 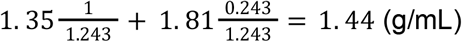 when *μ* increases from ~0h^-1^ to 0.7h^-1^. Oldewurthel et al. [6] found that up to these growth rates, the observed cellular dry mass density in *E. coli* is approximately constant at *ρ*_DM_=0.31g/mL. Accordingly, the empirical *ρ* decreases from 0.31 ÷ 1.39 = 0.223 to 0.31 ÷ 1.44 = 0.215, a 4% reduction.

The same set of experiments also showed that *ρ*_DM_ in *E. coli* decreases from 0.31g/mL to 0.28g/mL as *μ* increases further, from 0.7h^-1^ to 1.2h^-1^ [6]. Within this range of *μ*, *r* increases from 0.243 to 0.354 [1], and the overall density of cytosolic dry mass changes from 1.44g/mL to 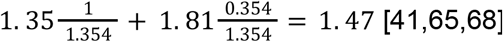. Accordingly, the empirical occupancy *ρ* decreases from 0.31 ÷ 1.44 = 0.215 to 0. 28 ÷ 1.47 = 0.191, a decrease of 13%.

## Availability of source code

The source code used for simulations in this study are uploaded to two GItHub repositories, one for the linear model and the other for the whole cell model:

1. https://github.com/TinPang/optimalCytosolicDensity_linearModel
2. https://github.com/TinPang/optimalCytosolicDensity_wholeCellModel

## Acknowledgements

We thank Deya Alzoubi and David Heckmann for providing the environments used in the ccFBA simulations. We thank Hugo Dourado and Deniz Sezer for helpful discussions. This work was supported by DFG Grants CRC 1310 to T.Y.P. and M.J.L. and by a grant of the Volkswagen Foundation in the “Life?” initiative to M.J.L..

## Author contributions

T.Y.P. developed the methodology and performed all analyses. M.J.L. and T.Y.P. conceived of the study, interpreted the results, and wrote the manuscript.

## Notes

### Competing Interest Statement

The authors have declared no competing interest.

### Summary of Updates

New analyses have been added to strengthen the arguments.

